# Methods matter: your measures of explicit and implicit processes in visuomotor adaptation affect your results

**DOI:** 10.1101/702290

**Authors:** Jana Maresch, Susen Werner, Opher Donchin

## Abstract

Visuomotor rotations are frequently used to study the different processes underlying motor adaptation. Explicit aiming strategies and implicit recalibration are two of these processes. Various methods, which differ in their underlying assumptions, have been used to dissociate the two processes. Direct methods, such as verbal reports, assume explicit knowledge to be verbalizable, where indirect methods, such as the exclusion, assume that explicit knowledge is controllable. The goal of this study was thus to directly compare verbal reporting with exclusion in two different conditions: during consistent reporting and during intermittent reporting. Our results show that our two conditions lead to a dissociation between the measures. In the consistent reporting group, all measures showed similar results. However, in the intermittent reporting group, verbal reporting showed more explicit re-aiming and less implicit adaptation than exclusion. Curiously, when exclusion was measured again, after the end of learning, the differences were no longer apparent. We suspect this may reflect selective decay in implicit adaptation, as has been reported previously. All told, our results clearly indicate that methods of measurement can affect the amount of explicit re-aiming and implicit adaptation that is measured. Since it has been previously shown that both explicit re-aiming and implicit adaptation have multiple components, discrepancies between these different methods may arise because different measures reflect different components.

## Introduction

Adaptation to visuomotor rotations is assumed to consist of at least two main underlying processes: an implicit process, termed here ‘implicit adaptation’ – which is slow, expressible at low reaction times, and rigid in different conditions – and an explicit process, termed ‘explicit re-aiming’ – which develops rapidly, requires a long preparation time, and is highly flexible (Taylor and Ivry 2011, Taylor, Krakauer et al. 2014, Bond and Taylor 2015, Haith, Huberdeau et al. 2015, Huberdeau, Krakauer et al. 2015, McDougle, Bond et al. 2015). Many different methods are used in visuomotor rotation tasks to assess these processes (Fernandez-Ruiz, Wong et al. 2011, Heuer and Hegele 2011, Taylor, Krakauer et al. 2014, Haith, Huberdeau et al. 2015, Huberdeau, Krakauer et al. 2015, McDougle, Bond et al. 2015, Werner, van Aken et al. 2015, Morehead, Taylor et al. 2017). These methods differ in their underlying assumptions: direct methods, such as asking subjects where they are aiming (Hegele and Heuer 2010, Taylor, Krakauer et al. 2014) and questionnaires at the end of the experiment (Hwang, Smith et al. 2006, Benson, Anguera et al. 2011) assume that knowledge is verbalizable; the former being more directed at subjects’ strategic corrections and the latter tapping on subjects’ general understanding of the perturbation. Indirect methods, such as manipulating subjects’ behavior using task design (Fernandez-Ruiz, Wong et al. 2011, Haith, Huberdeau et al. 2015, Morehead, Taylor et al. 2017), assume knowledge to be controllable.

Both types of methods have shortcomings, which are mainly addressed outside the motor field in consciousness research. Direct methods seem straightforward, but subjects may refrain from answering or be unable to report on certain experiences (Timmermans and Cleeremans 2015). Reports may also be contaminated by the observer paradox: asking subjects to produce subjective reports or to reflect on their own performance may influence the very processes that are being monitored (Newell and Shanks 2014, Timmermans and Cleeremans 2015). Specifically, subjects may thus develop more explicit knowledge about the task by being repeatedly reminded about it whereas implicit adaptation remains unaffected (Langsdorf, Maresch et al. 2019). Indirect methods may not confound measurement and awareness as fully as a direct measure, but there is still the danger that the experimental manipulation influences awareness. Additionally, indirect methods must be interpreted with care: since the subject has not declared explicitly what is in their head, we are reaching our conclusions through inference (Jacoby 1991, Cleeremans, Destrebecqz et al. 1998, Timmermans and Cleeremans 2015). Thus, there is no gold standard for measuring either implicit adaptation or explicit re-aiming. All measures must be viewed critically.

Visuomotor rotation experiments quite commonly rely on a direct measure of explicit re-aiming, the verbal report, and an indirect measure of implicit adaptation that we call exclusion. In the verbal report, subjects are asked to state out loud the intended aiming direction, usually based on landmarks presented visually around the target. Exclusion is measured in trials where subjects are told that the perturbation has been removed and asked to aim straight for the target (Hegele and Heuer 2010, Taylor and Ivry 2013). When measured after the adaptation, this is called aftereffect. The residual learned response in an exclusion trial is a measure of implicit adaptation.

The literature contains reports both of consistencies and inconsistencies between these measures (Taylor, Krakauer et al. 2014, Bond and Taylor 2015, Leow, Gunn et al. 2017, Bromberg, Donchin et al. 2019). The fact that they can be inconsistent dovetails with recent proposals that explicit re-aiming and implicit adaptation may both be composed of multiple components. For instance, McDougle et al. propose that explicit re-aiming may have different components that are computed and cached (McDougle and Taylor 2019). Others propose that implicit adaptation has labile and stable components (Miyamoto, Wang et al. 2014, Heuer and Hegele 2015, Kim, Parvin et al. 2019). One possibility is that different measures weight the components differently.

Thus, it may be a surprise that comparison of the measures has not yet been the focus of a dedicated, controlled study. Our goal is to compare explicit re-aiming and implicit adaptation under different experimental conditions as measured by verbal report and by exclusion. We hypothesize that differences we see in different measures across conditions may come from different expression of underlying components of explicit and implicit.

## Materials and methods

All data and scripts used in order to present the figures in this article are available in the online repository (https://osf.io/6yj3u/).

### Participants

Fifty-three right-handed participants completed the study and were randomly assigned to one of four groups: (1) consistent reporting (CR; n = 12; mean age: 26.5 [range 24-30]; 5 female), (2) intermittent reporting exclusion (IR-E; n = 17; mean age: 26.0 [22-34]; 7 female), (3) intermittent reporting inclusion (IR-I; n = 12; mean age: 24.4 [22-28]; 7 female) and (4) intermittent reporting exclusion and inclusion (IR-EI; n = 11; mean age: 25.3 [24-30]; 7 female). All subjects signed an informed consent form, which included basic information about their relevant medical status. Subjects were not included for participation if they had previously participated in visuomotor rotation research, if they had any neurological disorders or if they suffered from vertigo. Subjects were contacted and recruited through the department of Biomedical Engineering and received monetary compensation for their participation. The experimental protocol was approved by the Human Subjects Research committee of the Ben Gurion University, and followed the ethical guidelines of the university.

### Experimental apparatus and general procedures

Participants made center-out, horizontal reaching movements while holding on to the handle of a robotic manipulandum (Figure 1A). Movement trajectories were sampled at 200 Hz and a resolution of 0.3×10^−3^ degrees on each joint of the shoulder, which translates into a resolution in Cartesian coordinates of less than 0.2 mm. The stimuli were projected onto a horizontal plane in front of the participants (BenQ MS527, 3300 ANSI lumens), which also occluded vision of the subjects’ hand. At the beginning of each trial, the manipulandum guided the participants’ hand towards a white circle in the center of the display (5 mm radius), which was positioned approximately 45 cm away from the eyes of the subject. A small, circular cursor (white, 3 mm radius) provided continuous online visual feedback during each reach (except for trials in which no visual feedback was provided). After 50 ms, subjects were presented with a small target (red, 7 mm radius) at a distance of 10 cm from the origin. Targets could appear on a circle in one of eight possible, equally spaced locations (45° between targets: 0°, 45°, 90°, 135°, 180°, −135°, −90°, and −45°). Target locations were in a pseudorandom order such that all targets were experienced once before being repeated in the next cycle. Subjects were instructed to make fast and accurate shooting movements to the target, ‘slicing’ through the designated target. After leaving the starting position, participants had between 400 and 600 ms to reach the target in order for the target to turn green. This movement time was long enough for subjects to reach the target, however, it did not allow for corrections of movements. The target turned blue and yellow when movements were too slow or too fast, respectively. Additionally, a happy ding indicated target hit and an annoying buzz sound indicated target miss.

**Figure 1.**
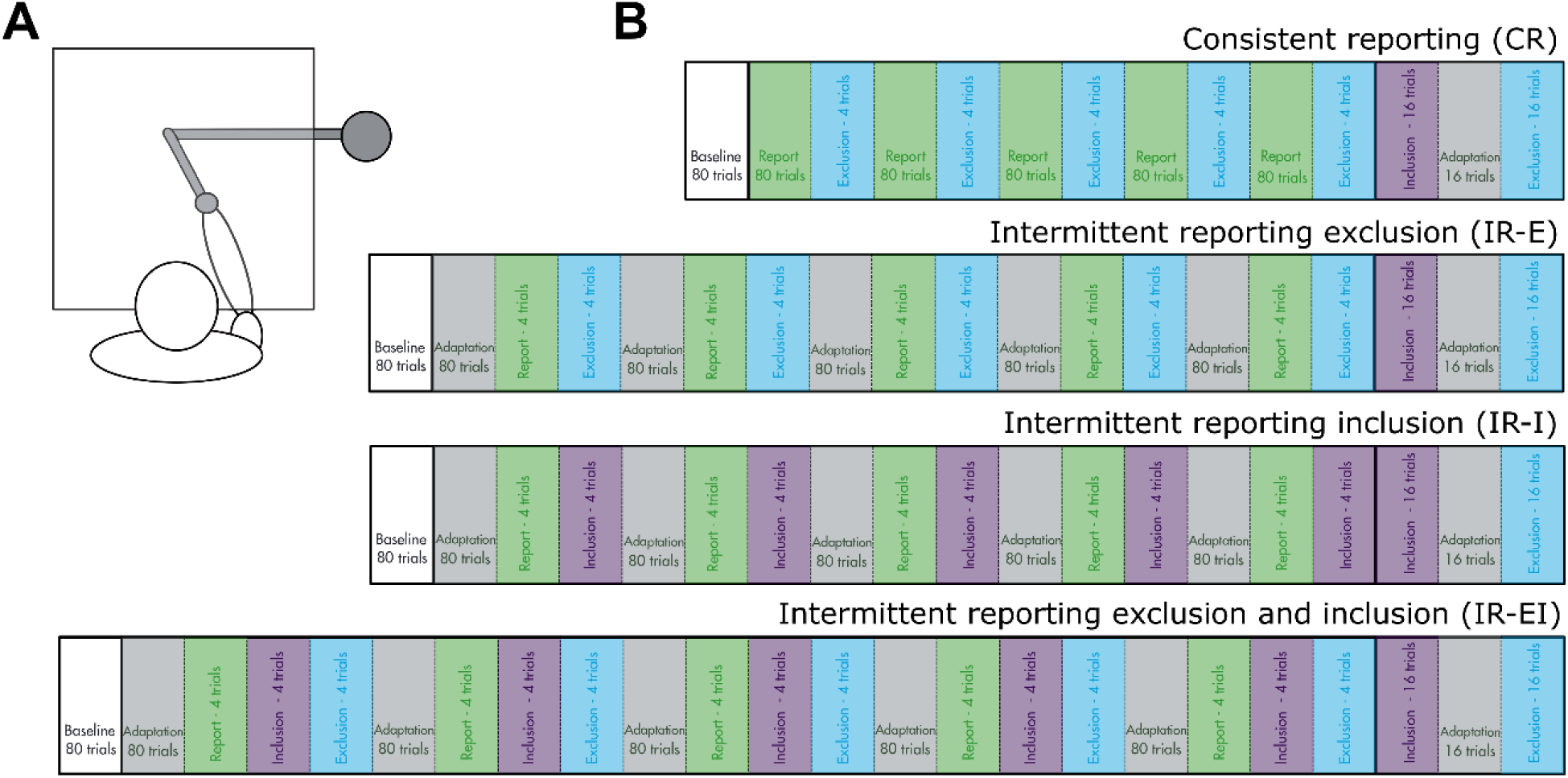
Experimental task. A) Subject were seated in front of a desk, which occluded sight of their arm and hand. With their right hand, subjects held the handle of a robotic manipulandum. The visual scene was projected onto the desk in front of the subjects. B) Procedures for the four experimental groups. Note that presentation of the different epochs is schematic and not to scale, i.e. the x-axis does not reflect the actual length of the individual epochs. Subjects in the CR group reported consistently throughout the entire adaptation (green boxes) while subjects in the IR-E, IR-I and the IR-EI groups performed regular adaptation blocks without reporting (gray boxes). At the end of each adaptation block, subjects performed four exclusion trials in the CR group (light blue boxes), report and exclusion trials in the IR-E group (green and light blue boxes), report and inclusion trials in the IR-I group (green and purple boxes) or report, exclusion and inclusion trials in the IR-EI group (green, light blue and purple boxes). Each group started with a baseline block and ended with a Process dissociation procedure block.

In some trials, subjects were asked to report their intended movement direction before beginning the movement. To this purpose, 42 landmarks, spaced 5.625° apart, were shown on a circle around the target. Landmarks were positive in the clockwise direction and negative in the counterclockwise direction (Figure 2A) and rotated with the target, such that the same landmarks would always appear in the same location relative to the target (Bond and Taylor 2015). Landmarks were only present during reporting trials. Verbal reports were recorded both online by the experimenter and digitally for later verification.

**Figure 2.**
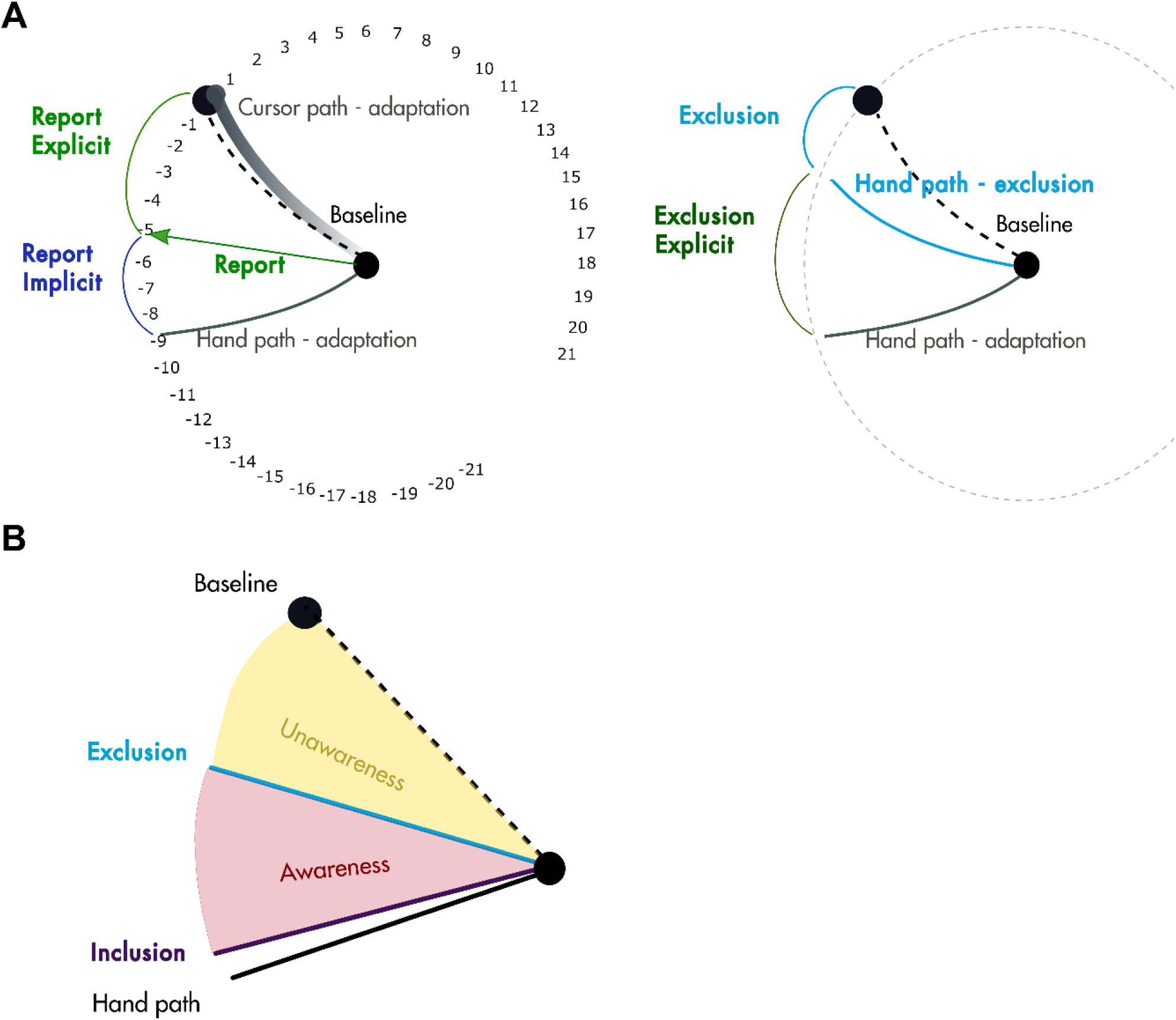
Intermittent and final measures. A) The difference between reported aiming directions and the hand path during adaptation provides us with the implicit_report_ measure (Taylor, Krakauer et al. 2014). B) The exclusion measure gives an indication of the implicit knowledge a subject has and thus cannot control. By subtracting the exclusion from the movement direction during the adaptation epoch (four trials), we obtain a measure of the explicit knowledge a subject had and can possibly control (explicit_excl_). Note that as opposed to the calculation in A), we here use the preceding adaptation epoch. **C)** Process dissociation procedure measure as adapted from Werner et al. (Werner, van Aken et al. 2015). The Awareness Index (AI, red) is derived from the difference between inclusion and exclusion whereas the Unawareness Index (UAI, yellow) is equal to the exclusion. AI and UAI may not add up to behavior (see Inclusion overshooting the hand path).

In each trial, the cursor and the hand movement were related in one of three different ways: veridical, no vision, or rotated. Veridical feedback was provided in most baseline trials. In some trials, the visual field was rotated so that the cursor was not precisely on the hand. This included a block of trials during the baseline designed to teach subjects how to report their aiming direction. In these trials, the visual field was rotated by between −30° and 30° on each trial by a different amount governed by a sum of sinusoids. We also used rotated trials in the adaptation blocks, where subjects were exposed to a consistent 60° rotation of the cursor. Finally, during the ‘refresh’ adaptation block between final exclusion and final inclusion, the cursor was rotated by 60°. There were two different types of no vision trials: inclusion trials and exclusion trials. The inclusion and exclusion task thus only differ in the instructions. Before inclusion, subjects were instructed to ‘use what was learned during learning’, and, before exclusion, subjects were asked to ‘refrain from using what was learned, and perform the movement as during baseline’. These instructions were identical to the ones used by Werner, van Aken et al. (2015) and Werner, Strueder et al. (2019). Instructions were given following a pre-determined instructions protocol, which was strictly (the full protocol is available on https://osf.io/6yj3u/). Prior to the experiment, each part of the experiment was shortly explained (familiarization, baseline, adaptation with intermittent test blocks, posttest). At this stage, subjects were told that it was important to remember what the baseline and adaptation blocks were since we would refer to these blocks in a later stage of the experiment (during both exclusion and inclusion trials in intermittent test blocks and in the post test). The names of the blocks were explicitly repeated at the beginning of both baseline and adaptation. Additionally, subjects were asked to restate the instructions in their own words before each intermittent and final test block so as to ascertain that they had understood the task. In order to ensure that our instructions would not reveal the nature of the perturbation, great care was taken in formulating the protocol not to use words like ‘re-aiming’ and ‘rotation’ or refer to dissociations between hand and cursor.

### Behavioral task

Prior to the start of the experiment, participants were told to expect a change in the task after the baseline block that may increase task difficulty. They were instructed to continue trying to slice through the target with the cursor. This was followed by a short familiarization session of 16 trials, during which subjects could ask questions and become familiar with the apparatus and task.

Familiarization was followed by a baseline block of 80 trials. The first half of the baseline block was 34 trials with veridical feedback interspersed with 6 trials with no visual feedback. Then, subjects conducted 40 trials of training in reporting their aiming directions (described above). The purpose of these trials was purely to train subjects in reporting aiming direction without exposing them to strong adaptation. Because of the changing perturbation, subjects were not able to either adapt or develop a working strategy.

After the baseline block, subjects did a series of 5 adaptation blocks of 80 movements each. In all adaptation blocks, visual feedback of the cursor was rotated by 60°. Subjects in the CR group reported their intended movement direction during the adaptation blocks.

Between each adaptation block, subjects did an intermittent measurement block. The composition of this block varied between the groups (see Figure 1B). The CR group did 4 exclusion trials. The IR-E group did four reporting trials and 4 exclusion trials. The IR-I group did four reporting and 4 inclusion trials. The IR-EI group did 4 reporting, 4 inclusion and 4 exclusion trials. Pilots using different sequences of the intermittent test blocks showed no order-effect and were thus always the same. As described above, instructions were shortly repeated before each intermittent measurement block. This explanation lasted about 20-30 s, whereas the timeframe between each individual measurement block was kept as short as possible, lasting approximately 5-10 s.

At the end of adaptation, subjects in all groups performed a posttest, comprised of the full process dissociation procedure. During a short break between the end of adaptation and the posttest, subjects were briefed on the last part of the experiment, reminding them of the names of the blocks and the inclusion and exclusion instructions. Depending on the group, additional care was taken to explain the type of trials this group had not experienced yet (e.g. inclusion for the IR-E group). The process dissociation procedure consisted of 48 trials, 16 of which were exclusion trials, 16 were ‘refresh’ adaptation trials, and the last 16 were inclusion trials. The order of exclusion and inclusion was counterbalanced between participants as suggested by Werner, van Aken et al. (2015).

After completion of the experiment, subjects filled out an online questionnaire about the experiment and the task. In the first part subjects had to answer open questions about the nature of the perturbation, and in the second part, the same questions were posed as multiple choice.

### Movement analysis

We characterized movement direction on each trial using the direction of the hand movement on that trial. Movement onset was defined as the time at which hand speed first exceeded 5cm/s. To assess movement direction, we tested the three different measures of movement direction: error at 8 cm, first maximal error (FME) and error at peak velocity (PVE). Error at 8 cm is the difference between the target angle and the average heading angle at 8 cm along the trajectory. FME is the angle between a line connecting starting and target dot and a line between movement onset and movement position at 100 ms after movement onset. Finally, PVE is the angle between a line connecting starting and target dot and a line between movement onset and movement position at peak velocity. We found large overlap between the different metrics, indicating that our results were not sensitive to the precise measure of movement direction we used and that subjects followed our instructions to produce fast shooting movements, which led to straight hand trajectories. We decided to use PVE as our measure for movement direction because it had the smallest within subject variability in our data. Average time until peak velocity was comparable across groups and the different types of trials (overall mean: 173 ms). All trajectories were rotated to a common axis with the target location at 0°. Reaction time for all groups was defined as the time between target appearance and movement onset. A trial was omitted if reaction times exceeded 1000 ms or if the subject’s movement fell short of reaching the target; 1.5 % of all trials were thus excluded. Please note that movement times were long enough for subjects to reach and ‘slice through’ the target on most trials. To visualize movement direction, we used bins of four trials each, which was the size of the intermittent measurement blocks.

In addition to movement direction, we had subject reports for intermittent reporting trials in all groups and also for the adaptation trials in the CR group.

We further summarized subject performance by calculating a number of additional measures from the movement directions and reporting.

#### Implicit_report_

the implicit_report_ was the difference between report explicit and movement direction (Figure 2A). While the report explicit provides us with an estimate of the explicit aiming subjects can verbalize, the implicit_report_ measures adaptation outside explicit verbal access.

#### Explicit_excl_

explicit_excl_ is the difference between movement direction at the end of adaptation immediately preceding the exclusion epoch and the movement direction in exclusion. This is the change in the movement direction caused by the subject’s re-aiming. As such, it measures the subject’s explicit control over their own behavior.

#### Awareness index

awareness index (AI) was calculated for each subject using average movement direction of the exclusion (*M*_exclusion_) and inclusion (*M*_inclusion_) epochs during the intermittent test (for the IR-EI group) or the posttest (for all groups) according to following formula:

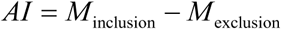

This calculation is based on the calculation used by Werner et al (2015, 2019) without the normalization they used to allow comparison of different rotation sizes. Without the normalization, the unawareness index in that paper reduces to the mean of the final exclusion. Thus, we do not calculate the unawareness index.

Note that we have two measures of explicit re-aiming based on the exclusion: explicit_excl_ (adaptation minus exclusion) and the awareness index (inclusion minus exclusion). Generally, the awareness index will indicate less explicit re-aiming than the explicit_excl_. The gap between them is a result of inclusion being generally lower than adaptation, i.e. closer to baseline (Werner, van Aken et al. 2015, Werner, Strueder et al. 2019). This may be because of the lack of visual feedback in the inclusion blocks or for other reasons.

### Statistical analyses

We used a Bayesian statistical approach very similar to the approach used in Werner, Strueder et al. (2019) to fit a linear model to our data. The Bayesian approach combines prior information about population parameters with evidence contained in the sample itself in order to create posterior probability distributions of the parameters (Kruschke 2014). In contrast to frequentist approaches, Bayesian probabilities are thus statements about probabilities of these parameters in general, and not about the specific sample in the study. Full details of the model are available in the online repository (https://osf.io/6yj3u/) and here we describe only the key points. The dependent variable for our linear model was the mean movement directions of each subject in each trial. The model’s dependent variables included the epoch type, the subject’s group, and the subject. Epoch types are defined below. The model also included the group by epoch type interaction and the subject by epoch type interaction. There were eight epoch types, see Table 1 for the overview of these epochs. Since not all groups did all epochs, we used imputation as described in Kruschke (2014), Ch. 20, to balance the model. Bayesian imputation requires a hierarchical model so that the statistics of the imputed data can be learned from the existing data. Thus, we made the group by episode interaction a hierarchical parameter. We gave it a t-distribution in order to allow outliers and sampled both the variance and the degrees of freedom of this distribution, as well as sampling the specific values for every episode and every group. The other coefficients of the model were assumed to be normally distributed and were drawn from a broad, uninformative prior. The standard deviation of movements around the linear model was assumed to differ among subjects and it was also sampled for each subject from a gamma distribution which was determined through hierarchical sampling. MATLAB code and model are available in the online repository (https://osf.io/6yj3u/).

**Table 1.**

| Epoch types | Number of epochs | Number of trials |
| --- | --- | --- |
| Baseline | 1 | 80 |
| Adaptation | 100 | 400 (5x80) |
| Report | 5 | 20 (5x4) |
| Exclusion | 5 | 20 (5x4) |
| Inclusion | 5 | 20 (5x4) |
| Final exclusion | 1 | 16 |
| Final inclusion | 1 | 16 |

The joint posterior distribution of the model’s parameters were sampled using JAGS (4.2.0, http://mcmc-jags.sourceforge.net/) called from MATLAB (2019b, the Mathworks, Natick, MA) using *matjags* (http://psiexp.ss.uci.edu/research/programs_data/jags/). We used 4 chains, 2000 burn in samples and 10000 samples per chain. Using standard diagnostics described by Kruschke (2010), we ensured that the chains had converged to a unimodal distribution for all parameters and that the results were consistent across chains. Again, full details of the sampling, the posterior samples and code to produce all the diagnostic plots are available in the online repository (https://osf.io/6yj3u/).

Using the sampled estimates of the linear coefficients, we calculated the posterior distribution of the mean movement direction for each epoch, subject and group. We report our results using the mean and 95 % high density interval (HDI) of these estimates of the mean. The HDI contains 95 % of the distribution, in which every point has higher credibility than any point outside this range. Furthermore, for group comparisons we specified a *region of practical equivalence* (ROPE) of −3° through 3° around a value of 0° difference between the groups. The limits of this ROPE are an arbitrary choice, which is based on what we consider a meaningful difference. We report all actual differences in our results, each reader can thus decide for himself what a meaningful difference may be. We report the percentage of the HDI that lies within the ROPE as a measure of the probability that the values are equivalent.

Unless otherwise indicated, brackets represent 95 % HDI.

## Results

### Adaptation and learning rates

The time course of movement directions showed a stereotypical learning curve for all four groups (Figure 3A). However, the learning curves differed between the CR and the intermittent groups: adaptation proceeded slower and reached asymptote later in the intermittent groups than in the CR group (see difference between blue learning curve and the rest). We compared differences in movement direction early and late in adaptation by computing our estimate of the mean over 1) the last eight trials of the first adaptation block and 2) the last eight trials of the last adaptation block (asymptote of learning). Figure 3B depicts the mean movement direction for early and late adaptation: the CR group showed greater change in movement direction during early adaptation (50.7° [48 – 53]) as compared to the IR-E (39.9° [38 – 42]), IR-I (39.4° [37 – 42]) and IR-EI group (40.3° [38 – 43]). These differences of 10°-11° are consistent across groups. Although the magnitude of the difference in movement directions between groups decreased to about 9°, it persisted until the end of adaptation (CR: 61.1° [59 – 34]; IR-E: 52.4° [50 – 54]; IR-I: 52.5° [50 – 55]; IR-EI 52.7° [50 – 55]).

**Figure 3.**
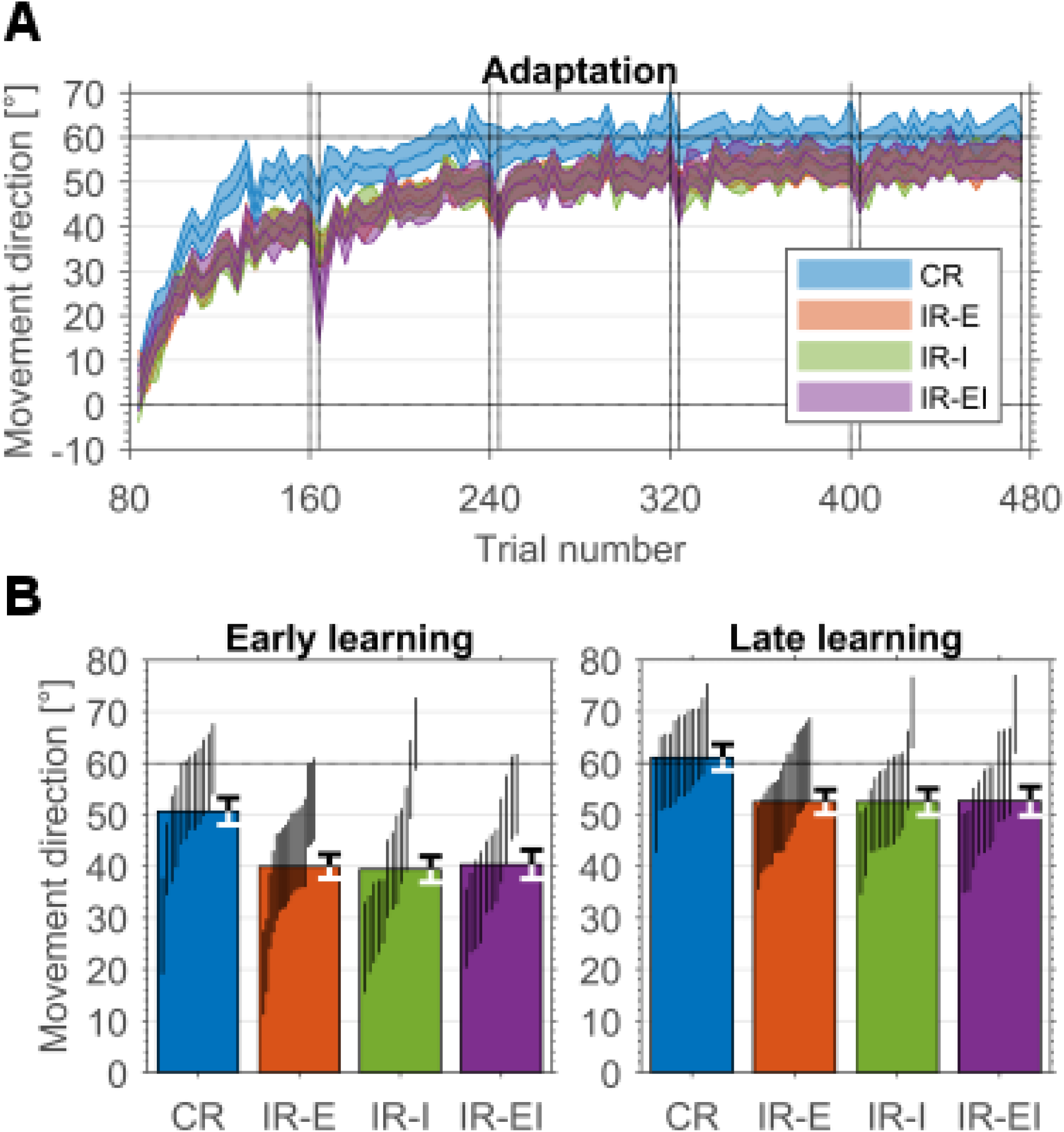
A) Binned estimated mean movement direction per group. Dashed line shows the end of baseline, dotted lines throughout adaptation show location of test blocks, note however, that the actual intermittent test are not presented here. The posttest (Process dissociation procedure) is not shown. B) Early and late movement direction. For the early adaptation, we used the last two epochs of the first adaptation block (8 trials) and, for the late adaptation, we used the last two epochs of the last adaptation block (8 trials). Vertical lines denote 95% HDI of mean of individual subjects. Error bars show 95% HDI of mean of the group.

### Intermittent measures

In Figure 4, we show the results of the intermittent test blocks. Results for these measures were comparable within the IR groups, we thus first present the IR-E group’s results as representative for all IR groups. Figure 4A depicts the time course of the intermittent measures for the CR and the IR-E group. Measures of explicit re-aiming are in green (report 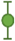 and explicit_excl_ 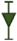) and measures of implicit adaptation are in blue (implicit_report_ 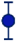 and exclusion 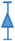). All measures in all groups changed over the first two epochs and then stabilized. After examining the data, we decided to use the average value of the last three epochs for any comparisons between measures and groups, as behavior had stabilized and subjects showed little variation between epochs.

**Figure 4.**
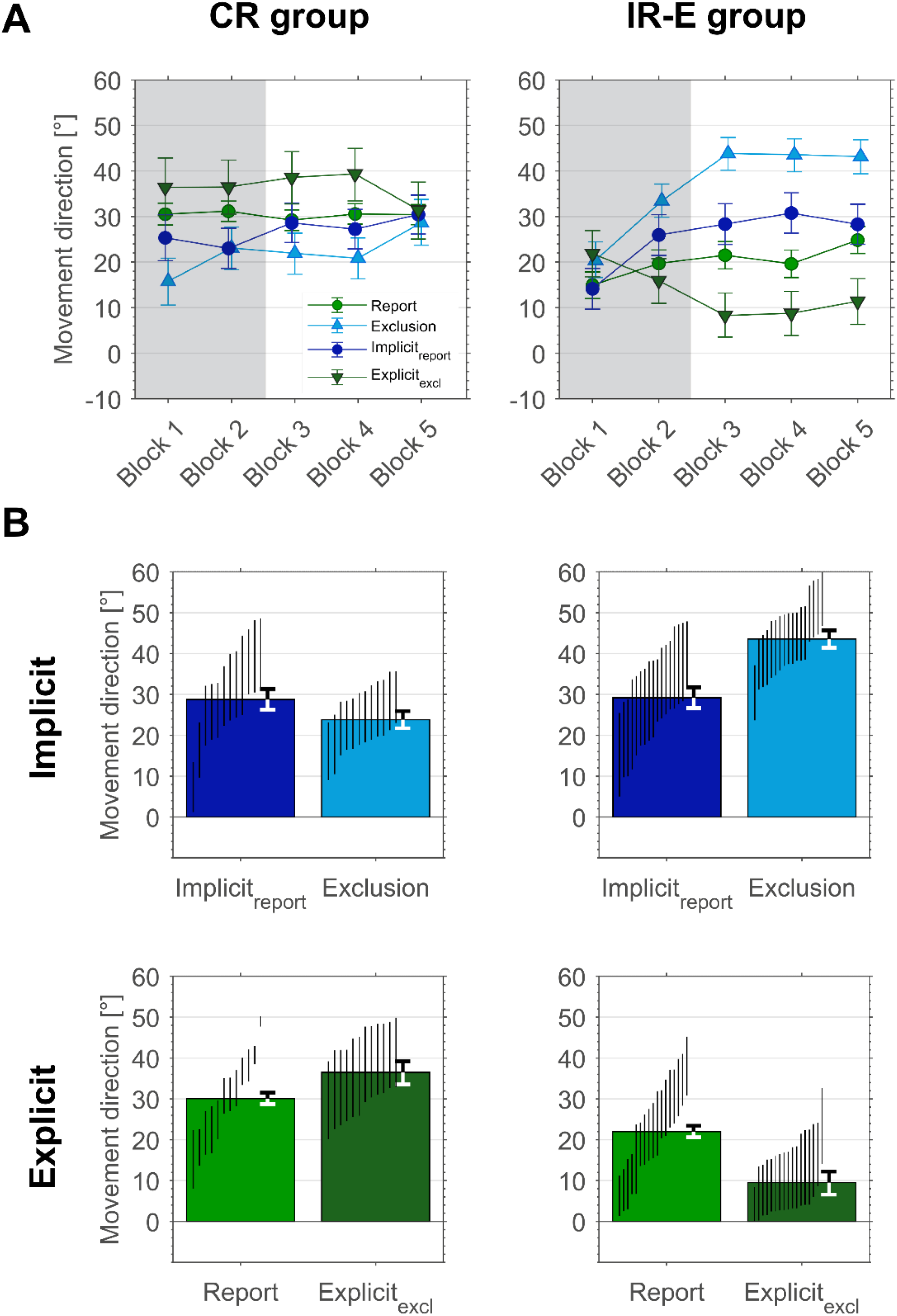
Intermittent explicit and implicit measures for the CR and IR-E groups. Intermittent measures were comparable across the IR groups, thus only measures for the IR-E group are presented here. Note that individual subject HDIs are sorted and do thus not correspond to subjects across sub-figures. A) Time course of report and exclusion measures per group. On the left we present the CR group, on the right the IR-E group. The grey shaded area denotes the first two intermittent test blocks. B) Intermittent measures per group. Bars show estimate of the mean of the last three test blocks (unshaded area in A). Top row shows implicit measures; bottom row shows explicit measures. Black lines depict 95% HDI for means of individual subjects; error bars show 95% HDI for means per group.

Implicit_report_ showed slightly higher levels of implicit adaptation than exclusion in the CR group (Figure 4B, top left column; 28.8° [26 – 31] and 23.8° [21 – 26], respectively). In contrast, implicit_report_ was much lower than exclusion in the IR-E group (Figure 4B, top right column; 29.1° [26 – 32] and 43.4° [41 – 46], respectively). The fact that implicit_report_ and exclusion are more different in the IR-E group arises because exclusion is different between the groups: implicit_report_ shows similar values between groups whereas exclusion values for subjects in the IR-E group are ∼ 20° higher than in the CR group.

Differences in measures of explicit re-aiming reflected our findings for the implicit adaptation. Reported aiming directions in the last three blocks were lower than the explicit_excl_ in the CR group (Figure 4B, bottom left; 30.1° [29 – 31] and 37.0° [33 – 40]). In contrast, the IR-E group showed much higher explicit re-aiming when using reported aiming directions than when using explicit_excl_ in (Figure 4B, bottom right; 22.0° [20 – 24] and 10.0° [7 – 12]). While implicit_report_ was similar between the groups, explicit re-aiming was greater in the CR group than the IR-E group according to both measures of explicit re-aiming. This difference was, however, much more striking for the explicit_excl_ than for the report.

In Figure 5 we present results also from the IR-EI group and show that our results are consistent across subjects and groups. The two measures produce different results, and these results are different for subjects in CR and IR groups. Exclusion is larger than implicit_report_ for all subjects in the IR groups (orange and purple) while exclusion is smaller than implicit_report_ for all subjects in the CR group (blue). Similarly, report is greater than explicit_excl_ in the IR groups and the opposite is true in the CR group. Taken together, our results indicate that reporting and exclusion are not measuring the same thing, although the difference may be hidden in some experimental conditions.

**Figure 5.**
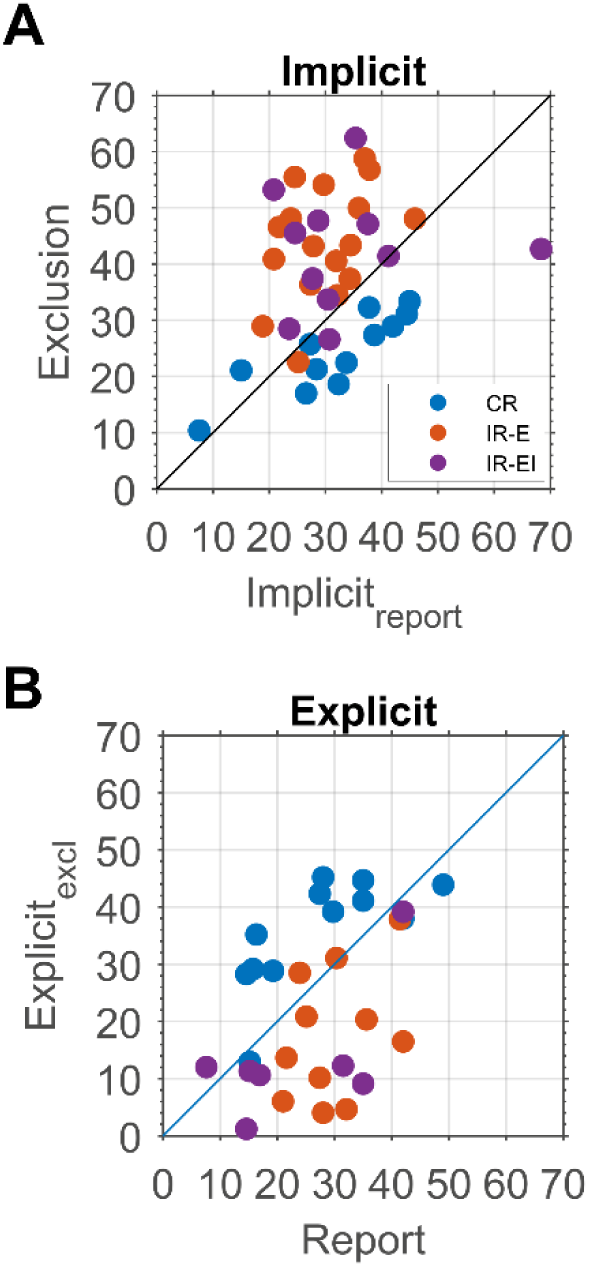
Explicit and implicit measures for individual subjects across all groups. A) Exclusion vs report implicit and B) exclusion explicit vs report.

### Intermittent measures IR-I and IR-EI group

Subjects in both the IR-I and IR-EI group performed inclusion as well as intermittent report trials. The IR-EI group, moreover, completed the entire process dissociation procedure in each intermittent measurement block. During inclusion, we ask subjects to perform the same movement as during adaptation, but without visual feedback. Figure 6A shows that subjects in both groups tended to undershoot during these trials, meaning less adaptation during inclusion trials than during the preceding adaptation trials (47.5° [45 – 50] and 43.4° [41 – 46] during inclusion for the IR-I ad IR-EI groups, respectively). The difference between adaptation and inclusion trials is relatively high, with the HDI of this difference not including zero for either IR-I group (5.8° [3 – 9]) or the IR-EI group (10.4° [7 – 14]). While the HDI and the ROPE of the IR-EI group do overlap, this overlap reflects less than 0.01% of the posterior density. The percentage of the posterior density in the overlap is slightly higher in the IR-I group (0.04%). Thus the two values are certainly not equivalent and may well be different for both groups. The fact that inclusion may not be the same as movement direction at the end of adaptation means that we need to be careful when interpreting the explicit_excl_.

**Figure 6.**
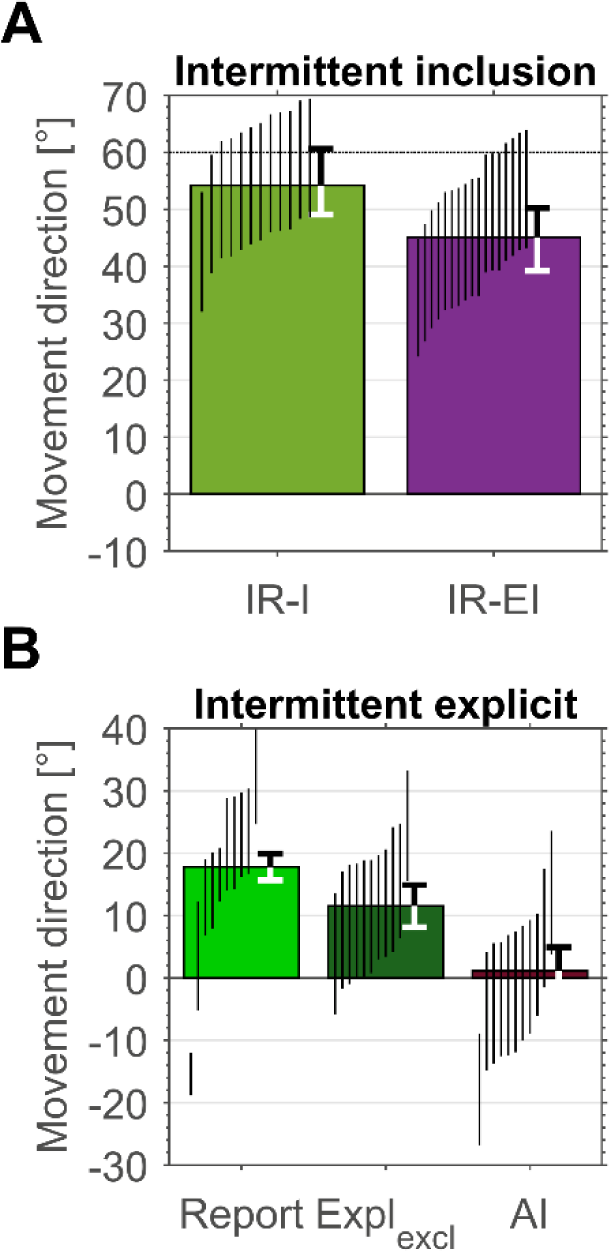
Intermittent measures for the IR-I and IR-EI groups. A) Intermittent inclusion for the IR-I and IR-EI group. B) Explicit intermittent measures for the IR-EI group. Shown is the mean the reported aiming direction, the mean exclusion explicit and the AI (calculated from the difference between inclusion and exclusion). Black lines denote 95% HDIs for individual subjects, error bars show 95% HDIs for the respective group.

We therefore calculated the AI using the intermittent measurement blocks of the IR-EI group. This intermittent AI enabled us to compare the two explicit measures we used with the additional AI with each other within the same group. As with the other intermittent measures, we used the last three intermittent measurement blocks for this comparison (described in the methods). Figure 6B shows that the AI estimates explicit re-aiming lower than both other measures, while the report provides the highest estimate (report: 17.8° [16 – 20]; explicit_excl_: 12.0° [8 – 15]; AI: 1.2° [-5 – 2]).

This further emphasizes that report estimates explicit re-aiming higher than exclusion based measures.

### Posttest

Our posttest, the process dissociation procedure, consisted of inclusion and exclusion blocks at the end of adaptation. At this point, each group received a short instruction about the posttest, which led to a short delay of approx. +/-2 min. Since subjects had experienced exclusion and/or inclusion intermittently (depending on the group), we expected these measures to produce similar results in the final test. However, this was not the case. In Figure 7, we show the different final measures per group. Notably, final exclusion is similar across groups (Figure 7A and B; CR: 31.6° [29 – 34], IR-E: 32.6° [31 – 35], IR-I: 37.6° [35 – 40] and IR-EI: 31.9° [29 – 34]), contrasting with our findings in the intermittent measurement blocks. This difference between intermittent and final exclusion was positive for the CR group (8.5° [5 – 11]) while it was negative for the IR-E (−11.7° [-15 – −9]) and IR-EI group (−9.7° [-14 – −6]). The consistency of this result across individuals is shown in Figure 7C, left scatter. In the discussion, we consider the possibility that this difference is the result of decay in a labile component of implicit adaptation caused by the time delay between the end of adaptation and the posttest (Hadjiosif and Smith 2013, Miyamoto, Wang et al. 2014, Morehead 2018, Kim, Parvin et al. 2019).

**Figure 7.**
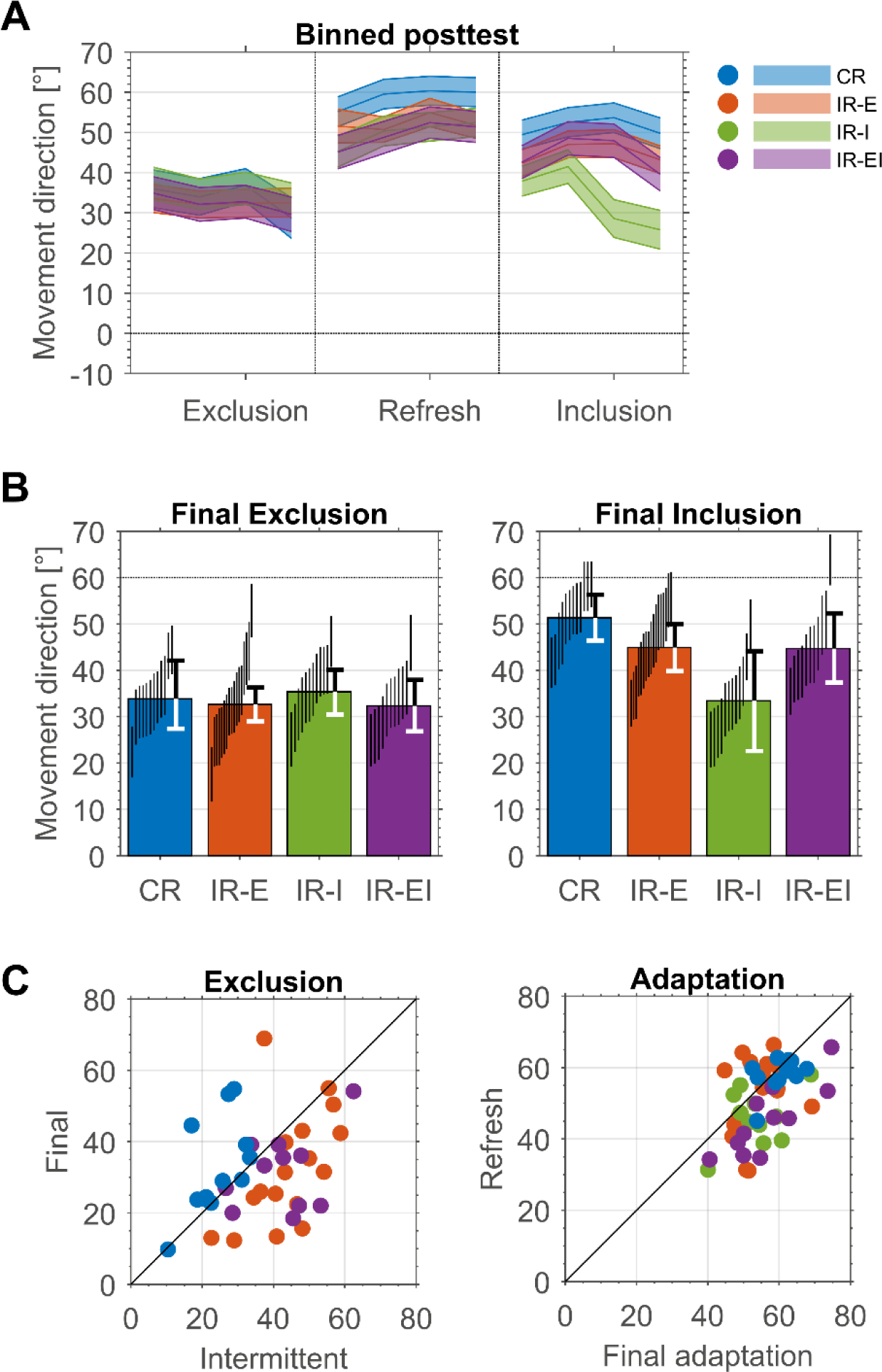
Posttest per group. Shown are the posttest measures and the differences between intermittent and final epochs. A) Movement directions during posttest. Note that exclusion is presented before inclusion, whereas the order was randomized across subjects. Shaded area denotes HDI. B) Final exclusion (left) and final Inclusion (right). Shown is the mean per 16 trials. Vertical lines represent 95% HDI for individual subjects, error bars show 95% HDI per group. C) Differences between intermittent and final movement directions for all groups. Left: Scatter plot of intermittent vs. final exclusion. Right: Scatter plot of movement direction at the end of adaptation vs. during refresh.

For this reason, we also considered the differences between the movement directions at the end of adaptation and in the posttest refresh block. We show in Figure 7 that movement direction generally reverts back towards baseline for most subjects between the end of adaptation and the refresh phase (Figure 7C, right scatter plot). This means that there is a decay in the learned adaptation during the interval between the end of adaptation and the beginning of the posttest. This is larger in the IR groups and it is consistent with the earlier hypothesis of a labile implicit component that is missing in the CR group.

Unlike final exclusion, final inclusion measures are different between groups. Subjects in the IR-I group showed the lowest values for the final inclusion (Figure 7B; IR-I: 31.7° [29 – 34]). Subjects in the IR-E and IR-EI group showed similar and higher inclusion values (IR-E: 46.2° [44 – 48]; IR-EI: 45.7° [43 – 48]), whereas subjects in the CR group were closest to full compensation during their inclusion (Figure 7B; CR: 51.0° [49 – 53]).]

Finally, our results for the AI reflect our findings described in the previous paragraph. The AI differed between groups and showed the same pattern as the final inclusion where subjects in the CR, in the IR-E and IR-EI groups showed more awareness than subjects in the IR-I (Figure 8; CR: 19.0° [16 – 22], IR-E: 13.4° [11 – 16] and IR-EI: 13.5° [10 – 17] as compared to IR-I: −5.0° [-8 – −1]).

**Figure 8.**
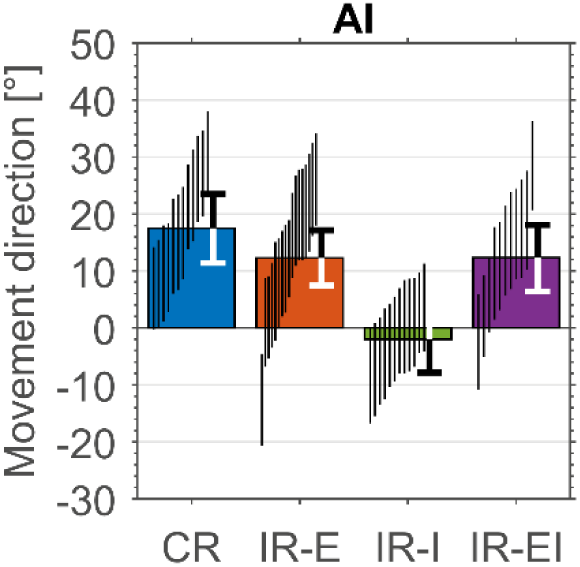
Awareness index for all groups calculated as the difference between final inclusion and exclusion. Vertical lines represent 95% HDI for individual subjects, error bars show 95% HDI per group.

### Reaction times

Figure 9 shows reaction times for the different groups during early (empty circles) and late adaptation (full circles) against our measures of explicit and implicit adaption. The CR-group showed consistently higher reaction times compared to the other groups. This is not surprising as subjects in this group were required to pause and state their intended aiming direction before moving towards the target (see also (Taylor, Krakauer et al. 2014, Bond and Taylor 2015, McDougle, Bond et al. 2015)). Reaction times did not differ between early and late adaptation and they were also not related to the different measures within each group (data not shown). The figure indicates that one possible explanation for the differences between the groups may be the differences in reaction times.

**Figure 9.**
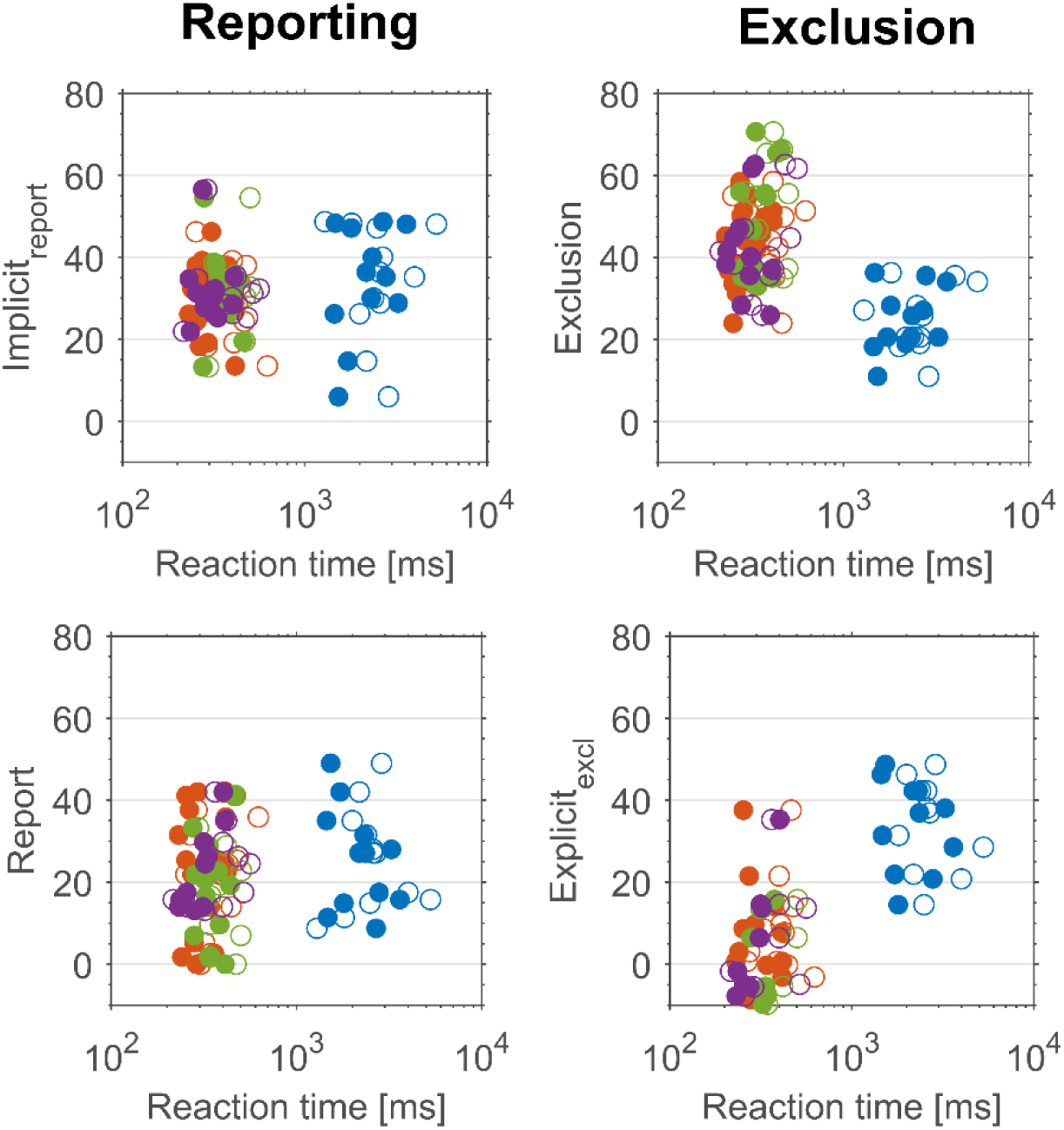
Reaction times for all groups. Shown are reaction times per measure (left and right column) for implicit and explicit processes (top and bottom row). Filled circles show reaction times in late adaptation (last two adaptation bins), whereas open circles show reaction times in early adaptation (first two adaptation bins). Note that the x-axis is presented on a logarithmic scale.

### Questionnaire

As described in the methods, each subject filled out an online questionnaire starting with three open questions about their understanding of the task. After this, the same questions were framed as multiple-choice questions. Answers and corresponding codes can be found in the online repository (https://osf.io/6yj3u/). We coded the answers to the open questions in categories ranging from 1 - 5 (1 means full understanding of the task, 5 is no understanding; see supplementary information). Three researchers separately assigned the codes to the answers of each subject. We present the median values and interquartile range of the codes across the three researchers and subjects per group. Subjects in the CR, IR-E and IR-EI group scored 2 points (2 ± 0, 2 ± 0.5, 2 ± 0.5 for CR, IR-E and IR-EI, respectively) in the open questions (subjects understood that a consistent rotation was applied), whereas the IR-I group scored 3 ± 0 points (subjects understood that a rotation or perturbation was applied but thought it was not consistent). This stands in line with our final inclusion and AI results where the IR-I group showed the lowest values.

In the multiple-choice questions, subjects were 1) asked to provide an explanation for the mismatch between the cursor and the hand and 2) asked in which direction the cursor was rotated relative to their hand. Subjects in the CR group were able to better characterize the mismatch than subjects in the IR groups (percentage correct: 92%, 53%, 67% and 75% for the CR, IR-E, IR-I and IR-EI group, respectively). When asked to classify the direction of the rotation, subjects in all groups were close to chance level (percentage correct: 58%, 41%, 58% and 42% for the CR, IR-E, IR-I and IR-EI group, respectively). Here, the first multiple choice question reflects a higher understanding if the task for subjects in the CR group, however, no further parallels can be drawn with our behavioral results.

## Discussion

Our goal was to compare measures of explicit re-aiming and implicit adaptation in a visuomotor rotation task to determine whether they measure the same thing. Our central result is that they do not. We compared two measures of explicit re-aiming (report and explicit_excl_) and of implicit adaptation (implicit_report_ and exclusion) in four conditions. We had one consistent reporting group (CR), which also performed intermittent exclusion trials, and three intermittent reporting groups who performed exclusion trials (IR-E), inclusion trials (IR-I) or both exclusion and inclusion trials (IR-EI). Performing both inclusion and exclusion trials provides a full process dissociation procedure as used by Werner et al. (2015, 2019), which provides an additional measure based on exclusion. Our measures produced consistent results in the CR group (Figure 5 bottom, blue dots), but in the other groups we found differences in both explicit re-aiming (report was greater than exclusion) and in implicit adaptation (exclusion was greater than report). This can be seen in Figure 5 in the orange and purple dots. The results of the process dissociation procedure from the IR-EI group suggest that this discrepancy in the IR groups may be even greater because considering exclusion without also considering inclusion may overestimate explicit_excl_ (Figure 6B).

It is worth making clear what these differences actually mean. A typical subject in the CR group will have fully adapted to the 60° rotation. When asked, however, they will report an aiming angle of only 30°. When asked to shift their aim back to the target, they will shift their aim by more than 30°. Typically, they will shift their aim by 40°, suggesting that they may have more control over their aiming direction than expressed in the report.

In contrast, a typical subject in one of the IR groups will have adapted partially. Rather than a full 60° adaptation, they will move towards 50°. When asked, they will report an aiming angle of 20°. However, when asked to shift their aim back to the target they will shift their aim by only 10°, showing that they have less control over their aiming direction than they think they do.

The differences in the explicit re-aiming shown by the two measures may reflect biased measurements. For instance, it is possible that report underestimates the explicit re-aiming in the CR group. The CR group has faster and more complete adaptation than the IR groups (Figure 3), and these have previously been associated with more explicit re-aiming (Benson, Anguera et al. 2011, Haith, Huberdeau et al. 2015, Leow, Gunn et al. 2017). This is in line with previous work suggesting that verbal responses may not reveal all of subjects’ knowledge, especially when knowledge is held with low confidence or is retrieved in a different context (Eriksen 1960, Nisbett and Wilson 1977, Shanks, Rowland et al. 2005). Conversely, it has been suggested that prediction tasks – such as reporting – during sequence-learning, concept-learning and grammar-learning are based on feelings of familiarity and, therefore, lead to an overestimation of awareness (Cleeremans and Elman 1993, Shanks and John 1994). Accordingly, explicit re-aiming as measured by report may be overestimating the actual explicit re-aiming as a result of feelings of familiarity. Since the CR group was reporting continuously, feelings of familiarity – and thus overestimation of explicit re-aiming – should be greater in this group as compared to the IR groups. Although the CR groups shows less variability in their reports – an indication that subjects in this group were more accustomed to reporting – our data shows the opposite: lower explicit re-aiming as measured by report than by explicit_excl_ in the CR group and larger explicit re-aiming as measured by report than by explicit_excl_ in the IR groups. Explicit_excl_ could also be biased. The change in exclusion between intermittent and final in the IR groups might reflect bias in the intermittent exclusion. Thus, none of our measures necessarily reflects a privileged measure of explicit re-aiming and all may have biases. However, across both measures and both levels of reporting, we do not see a theory of bias that would explain all of the differences in our results.

An alternative explanation for the dissociation between our two methods may be that explicit re-aiming is not a single process. This is not a new idea (Redding and Wallace 1996, Redding, Rossetti et al. 2005, Witt and Proffitt 2007). Specifically regarding visuomotor rotations and reaching movements, McDougle and Taylor (2019) theorized that explicit re-aiming is a combination of calculated and memorized re-aiming. Our results might arise because the CR and IR groups have different levels of each of these components and, in addition, different methods of measuring re-aiming reflect different combinations of these components. For instance, in the McDougle and Taylor model, calculated re-aiming is re-computed for each movement, a computation that takes time. Memorized re-aiming is not re-computed and does not require a longer reaction time, but it is limited to a small number of targets. Since our CR group had consistently longer reaction times than our IR groups, this may have contributed to greater calculated than cached learning in the CR group. Under this assumption, we generated the graph in Figure 10 that shows one (of several) ways that a specific sensitivity of report and exclusion to the different component could lead to the results we see. The specific model that generated this figure and associated calculations are available in the online repository (https://osf.io/6yj3u/). Multiple levels or components of re-aiming are one possible interpretation that could explain our results. However, much additional work is required to determine the actual underlying components and the way they are reflected in the different measures of re-aiming.

**Figure 10.**
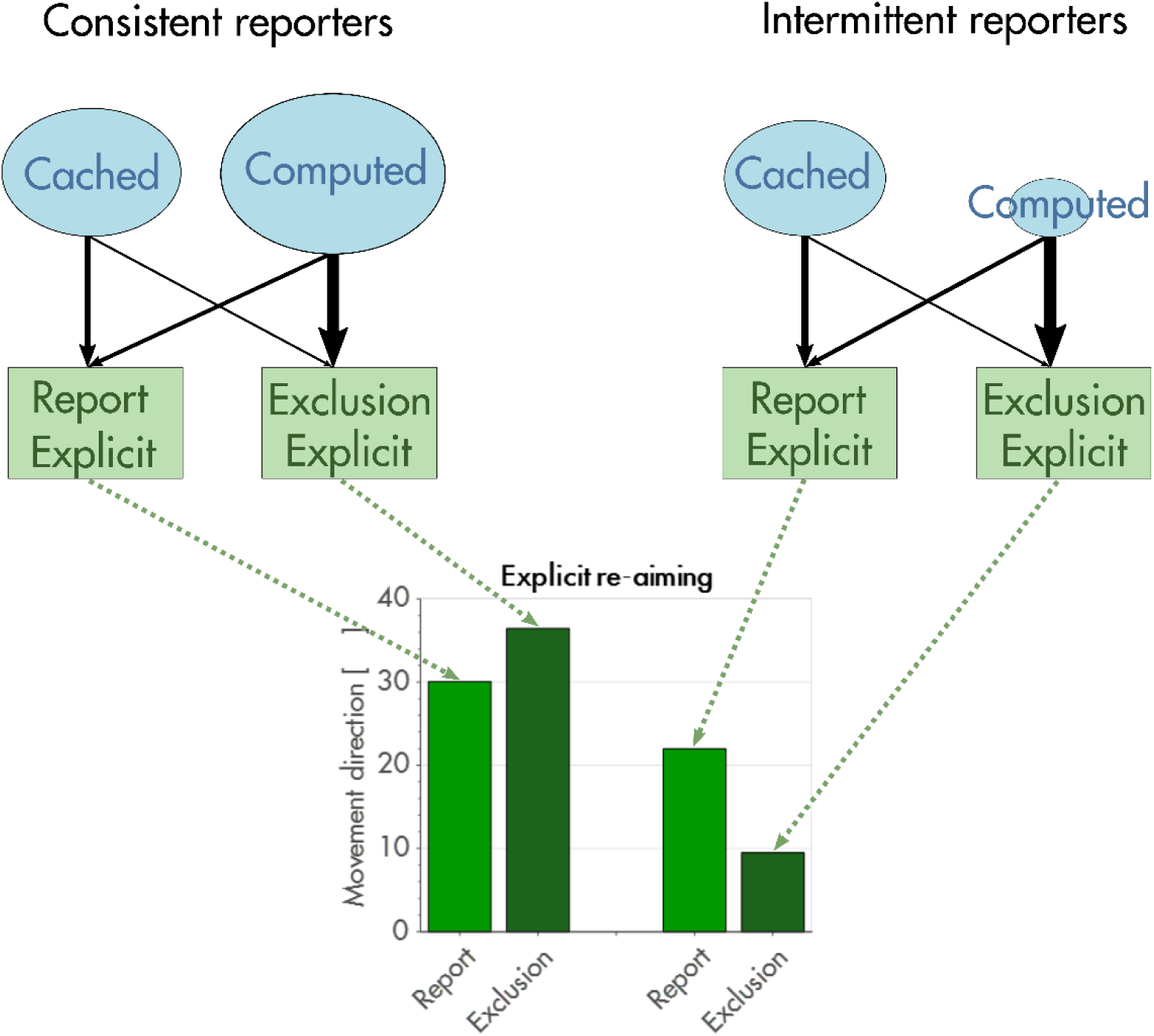
Example of how computed and cached components could combine in order to form explicit re-aiming as measured by exclusion and report for both consistent and intermittent reporters. Our model assumes that each component is weighted differently for each measure. One possible solution for the different components is the following: in the CR group, report reflects 37° cached and 28° computed re-aiming. In the IR groups, report reflects 5° cached and 27° computed re-aiming. Report would thus primarily reflect computed re-aiming (77%) and exclusion would primarily reflect cached re-aiming (77%).

Discrepancies in estimates of implicit adaptation measured by report and exclusion have been reported previously (Bond and Taylor 2015, Leow, Gunn et al. 2017). Leow et al. (2017) compared reporting and non-reporting groups. Their results are congruent with ours: the aftereffect shows more implicit knowledge in the non-reporting group than in the reporting group. One consideration that should be mentioned at this point are the longer delays due to reporting in the CR group, which may have led to larger decay of labile implicit in this group, and, possibly to more explicit re-aiming. However, reporting times were so short (2 – 4 s) that substantial decay of implicit adaptation seems unlikely. Bond et al. (2015) report some conditions where aftereffect showed less implicit knowledge than report. The direction of these effects is the opposite from our own, but the task conditions are also different from those in our experiments. Thus, different measures of implicit adaptation may produce different results, and these differences may depend on task conditions.

Differences in adaptation between reporting and non-reporting groups have been reported previously (Taylor, Krakauer et al. 2014, Leow, Gunn et al. 2017, Bromberg, Donchin et al. 2019). Actual asymptotic values differ between studies, which may be the result of different rotation sizes (we are using 60°, while most other studies use 30° or 45°). Larger rotation sizes may lead to more explicit re-aiming and, consequently, to a higher asymptote. However, it has also been proposed that the difference in asymptote does not arise from larger contributions of explicit re-aiming, but from the additional time subjects have when reporting, which allows for improved movement planning (Langsdorf, Maresch et al. 2019).

Some concerns regarding our data need to be addressed. Exclusion is large in the IR groups compared with previous findings regarding implicit adaptation in visuomotor rotation tasks (Hegele and Heuer 2010, Benson, Anguera et al. 2011, Heuer and Hegele 2011, Bond and Taylor 2015, Heuer and Hegele 2015, Morehead, Qasim et al. 2015, Leow, Gunn et al. 2017, McDougle, Bond et al. 2017). This could be a result of differences in task design: the use of a robotic manipulandum rather than a tablet, passive rather than active return of the hand, or the use of intermittent reporting. Interestingly, the large exclusion in the IR groups actually decreases over time, so that when it is measured several minutes after adaptation, as part of the process dissociation procedure, subjects in all groups have the same amount of exclusion (Figure 6B). Some have reported that implicit adaptation is composed of stable and labile components (Hadjiosif and Smith 2013, Miyamoto, Wang et al. 2014, McDougle, Bond et al. 2015, Morehead 2018), and it may be that intermittent reporting leads to more labile implicit, which, in part, explains the decay of the exclusion in the IR groups (Figure 6C). Since temporally labile implicit has been shown to decay at a time constant of 15-30 s (Hadjiosif and Smith 2013, Morehead 2018), one could postulate that our intermittent measures may have been affected by this decay. This may indeed be the case in the inclusion in the IR-EI measure, since this group performed inclusion more than 30sec after the reporting trials (also see Figure 6A: the IR-EI group shows less inclusion than the IR-I group). In all other groups, reporting was almost immediately (5-10 s) followed by exclusion or inclusion trials, rendering large decays improbable. Another consideration that may have led to differences between intermittent and final estimates is a change in subjects’ understanding of the instructions. However, since instructions were consistent across groups, we would expect the same change in understanding in all groups, which was not the case.

We also need to consider possibility that our exclusion measure is biased because it generalizes around the intended aiming direction rather than around the actual movement (Day, Roemmich et al. 2016, McDougle, Bond et al. 2017). Our results show slightly less exclusion than implicit_report_ in the CR group, which stands in line with generalization around the intended aiming direction.

However, our data shows a difference of ∼5° (Figure 4B top and Figure 4C) while generalization studies show ∼10° difference between exclusion and implicit_report_. Moreover, our IR groups show exactly the opposite effect of what generalization around the intended aiming direction would predict: exclusion is ∼15° larger than implicit_report_. It is also worth noting that our task conditions were quite different than those used in the generalization studies. Specifically, we used eight targets rather than one and we used online instead of endpoint feedback (Krakauer, Pine et al. 2000, Day, Roemmich et al. 2016, McDougle, Bond et al. 2017, Schween, Taylor et al. 2018). These differences could affect the width of the generalization function.

Finally, we want to revisit the starting point of this paper: the fact that all methods have their shortcomings. When we focus on quantifying subjective experience, all measures will face interpretational complications. As such, it has been previously suggested in memory research and sequence learning that subjects may fail to perform exclusion and inclusion reliably (Graf and Komatsu 1994, Dodson and Johnson 1996) – either as a result of not understanding the instructions or because of the complexity of the instructions – or they may use different strategies for exclusion and inclusion leading to distorted estimates of explicit and implicit processes (Barth, Stahl et al. 2019). Failure to exclude and/or include may then lead to biased estimates of the AI. Although great care was taken to make instructions as clear as possible (see methods: Experimental apparatus and general procedures) and although our exclusion and inclusion results largely match previous reports in visuomotor rotation studies (Werner, van Aken et al. 2015, Werner, Strueder et al. 2019), it is possible that our methods are not measuring exactly what we think they are. Our results (and results of others) might thus be plagued by systematic errors. The field needs more comparisons across measures, such as the ones in this paper, to make progress in overcoming this methodological gap.

In summary, our central finding is that different approaches to measuring explicit re-aiming and implicit adaptation lead to different results. Thus, they may reflect different components of the explicit and implicit processes. We speculate that explicit re-aiming is not a single process but made up of different components that contribute differently to measures of re-aiming depending on the task conditions. Methodological and theoretical developments are needed to fully understanding how these components combine to produce reaching movements. We need a new, updated model of the explicit and implicit processes during adaptation.

## Acknowledgements

We would like to thank Sivan Hazan for technical support and help with the data collection. Furthermore, we thank Dr. Shlomi Haar, Dr. Matthias Hegele, Dr. Liad Mudrik and Dr. Ronen Segev for comments and discussions about our data.

## Funding

This work is part of the PACE project (itn-pace.eu), which received funding from the European Union’s Horizon 2020 research and innovation program under the Marie Skłodowska-Curie grant agreement No 642961.

## Conflict of interest statement

No conflicts of interest, financial or otherwise, are declared by the authors.

## Author contributions

J.M., O.D. and S.W. contributed to the idea and design of the experiment. J.M. carried out the data collection, processed the experimental data, drafted the manuscript and designed the figures. O.D. performed the Bayesian statistical analysis. All authors discussed the results and contributed to the final manuscript.

## Data Accessibility statement

All data and MATLAB scripts used for analysis and plotting of the figures are available in the online repository (https://osf.io/6yj3u/).

## Abbreviations

CR: Consistent reporting
IR-E: Intermittent reporting exclusion
IR-I: Intermittent reporting inclusion
IR-EI: Intermittent reporting exclusion & inclusion
AI: awareness index
HDI: High density interval
ROPE: region of practical equivalence

## Notes

https://osf.io/6yj3u/

### Competing Interest Statement

The authors have declared no competing interest.

### Summary of Updates

This is the revised version of the original manuscript following comments of two reviewer’s and invitation to resubmit (after submission to EJN).

